# *gbpA* and *chiA* genes are not uniformly distributed amongst diverse *Vibrio cholerae*

**DOI:** 10.1101/2021.02.11.430729

**Authors:** Thea G. Fennell, Grace A. Blackwell, Nicholas R. Thomson, Matthew J. Dorman

## Abstract

Members of the bacterial genus *Vibrio* utilise chitin both as a metabolic substrate and a signal to activate natural competence. *Vibrio cholerae* is a bacterial enteric pathogen, sub-lineages of which can cause pandemic cholera. However, the chitin metabolic pathway in *V. cholerae* has been dissected using only a limited number of laboratory strains of this species. Here, we survey the complement of key chitin metabolism genes amongst 195 diverse *V. cholerae.* We show that the gene encoding GbpA, known to be an important colonisation and virulence factor in pandemic isolates, is not ubiquitous amongst *V. cholerae.* We also identify a putatively novel chitinase, and present experimental evidence in support of its functionality. Our data indicate that the chitin metabolic pathway within the *V. cholerae* species is more complex than previously thought, and emphasise the importance of considering genes and functions in the context of a species in its entirety, rather than simply relying on traditional reference strains.

**Impact statement:** It is thought that the ability to metabolise chitin is ubiquitous amongst *Vibrio* spp., and that this enables these species to survive in aqueous and estuarine environmental contexts. Although chitin metabolism pathways have been detailed in several members of this genus, little is known about how these processes vary within a single *Vibrio* species. Here, we present the distribution of genes encoding key chitinase and chitin-binding proteins across diverse *Vibrio cholerae*, and show that our canonical understanding of this pathway in this species is challenged when isolates from non-pandemic *V. cholerae* lineages are considered alongside those linked to pandemics. Furthermore, we show that genes previously thought to be species core genes are not in fact ubiquitous, and we identify novel components of the chitin metabolic cascade in this species, and present functional validation for these observations.

**Data summary:** The authors confirm that all supporting data, code, and protocols have been provided within the article or through supplementary data files.

1. No whole-genome sequencing data were generated in this study. Accession numbers for the publicly-available sequences used for these analyses are listed in Supplementary Table 1, Table 2, and the Methods.
2. All other data which underpin the figures in this manuscript, including pangenome data matrices, modified and unmodified sequence alignments and phylogenetic trees, original images of gels and immunoblots, raw fluorescence data, amplicon sequencing reads, and the R code used to generate Figure 7, are available in Figshare: https://dx.doi.org/10.6084/m9.figshare.13169189 (Note for peer-review: Figshare DOI is inactive but will be activated upon publication, please use temporary URL https://figshare.com/s/7795a2d80c13f694f8fa for review).

## Introduction

The *Vibrio* genus of marine γ-proteobacteria contains a number of virulent human pathogens, of significant public health concern [1]. Most notorious of these pathogens is the *Vibrio cholerae* species, members of which are the aetiological agent of cholera in humans [2, 3]. Two biochemically-defined and distinct *V. cholerae* biotypes are associated with cholera pandemics. Classical biotype *V. cholerae* are believed to have caused the first six pandemics [2–4], whilst the current seventh pandemic (1961-present) is attributed to El Tor biotype *V. cholerae* [5, 6]. Genomic evidence has shown that classical *V. cholerae* form a discrete phylogenetic lineage from the lineage causing the seventh pandemic, dubbed the <u>s</u>eventh <u>p</u>andemic <u>E</u>l <u>T</u>or lineage (<u>7PET</u>) [7–12]. It is to these two pandemic lineages that commonly-used El Tor and classical biotype laboratory strains belong. Although cholera is estimated to cause 100,000 deaths annually worldwide [13], other *Vibrio* species can also cause enteric and extraintestinal disease in humans. For example, *Vibrio vulnificus* can cause septicaemia and systemic infection in humans [14], and *Vibrio parahaemolyticus* can cause gastrointestinal infection, septicaemia, and wound infections [1, 15]. Other Vibrios may be pathogenic to livestock and other animals, such as *Vibrio nigripulchritudino* which is a pathogen of farmed shrimp [16, 17], and *Vibrio anguillarum*, which causes vibriosis in multiple species of fish [18].

In spite of differences in the types of disease which *Vibrio* species may cause, there are several commonalities amongst members of this genus. For example, it has been suggested that the ability to grow on chitin is a ubiquitous phenotype amongst the *Vibrionaceae* [19], and therefore that all *Vibrio* species are capable of metabolising chitin, a highly-abundant polymer of *N*-acetyl-β-d-glucosamine (GlcNAc) [20]. This is directly relevant to the environmental lifestyles of Vibrios - for example, *V. vulnificus* colonises and grows on the surface of chitinaceous animals such as shellfish [21]. *V. parahaemolyticus* secretes a chitinase and can adsorb onto particulate chitin and copepods [22]. Similarly, *V. cholerae* can metabolise chitin [23], has chitinase activity and can adsorb on chitinous substrates [24], and can colonise chitinous surfaces such as those of copepods [25]. Chitin metabolism is linked to other aspects of *Vibrio* biology, including the regulation of natural competence [26–28], and to the survival of *V. cholerae* in the context of the intestine during an infection [29].

The pathways by which chitin is degraded and utilised by *V. cholerae* have been described in detail [30], as it has been in other members of the genus (e.g., [31–38]). Although a comprehensive review of the chitin utilisation pathway is beyond the scope of this manuscript, it is important to appreciate the complexity of this pathway. Chitin degradation, import, and metabolism in *V. cholerae* involves at least 27 proteins, 24 encoded by genes on chromosome 1, and three by genes on the smaller chromosome 2 [19]. Here, focus will be directed to the initial stages of chitin metabolism - adhesion to a chitinous substrate, and expression of extracellular degradative chitinase enzymes.

The first step in chitin metabolism is the attachment of *V. cholerae* to chitinaceous surfaces through interactions with *N*-acetyl-β-d-glucosamine (GlcNAc). This is mediated both by the mannose-sensitive haemagglutinin (MSHA) pilus and the chitin adhesin GbpA (encoded by *VCA_0811*, accession # AAF96709.1) [23, 39]. Although GbpA was initially identified as a putative chitinase enzyme [23], it was shown to be an adhesin induced by GlcNAc which enabled *V. cholerae* to attach to chitinous substrates [23]. Subsequently, it was found that as well as mediating attachment of *V. cholerae* to chitin, GbpA is also required for the successful colonisation of the intestine [39]. This is thought to occur through interactions with mucin - GbpA interacts with mucin in the intestine, and *gbpA* transcription increases upon exposure of *V. cholerae* to mucin [40]. The crystal structure and domain architecture of GbpA have been determined [41], and the fourth domain of GbpA is structurally similar to the chitin-binding domain of known chitinases [41]. Evidence also suggests that GbpA has lytic polysaccharide monooxygenase activity [42], and that GbpA activity is higher at low population densities due to the activity of quorum-sensing-regulated proteases [43].

Once *V. cholerae* adheres to a chitinaceous surface, extracellular endochitinase enzymes are required for the bacterium to hydrolyse complex chitin polymers into oligosaccharides which can be imported into the cell for further metabolism [34]. As many as seven putative endochitinases have been identified in *V. cholerae* [19, 44, 45], two of which, ChiA-1 (encoded by *VC_1952*, accession # AAF95100.1) and ChiA-2 (encoded by *VCA_0027*, accession # AAF95941.1), are the principal chitinases required for *V. cholerae* chitin catabolism [23, 30, 44, 46]. ChiA-1 was first shown to be an extracellular chitinase in 1998 [47]; subsequently, ChiA-2 was shown to be important for intestinal colonisation and for metabolising mucin in the intestine by *V. cholerae* strain N16961 (N16961) [29]. ChiA-2 is also the most highly-expressed chitinase in El Tor biotype *V. cholerae* strain E7946 [44]. Both ChiA-1 and ChiA-2 are essential for *V. cholerae* to grow in media supplemented with colloidal chitin [23]. Once chitin oligomers have been digested by extracellular chitinases, the resultant oligosaccharides are thought to enter the bacterial periplasm *via* the chitoporin ChiP (encoded by *VC_0972*, accession # AAF94134.1) and by other as-yet-uncharacterised porins [23, 35, 48], and subsequently transported to the cytoplasm *via* PTS and ABC-type transporters [19, 35] (Figure 1).

**Figure 1.**
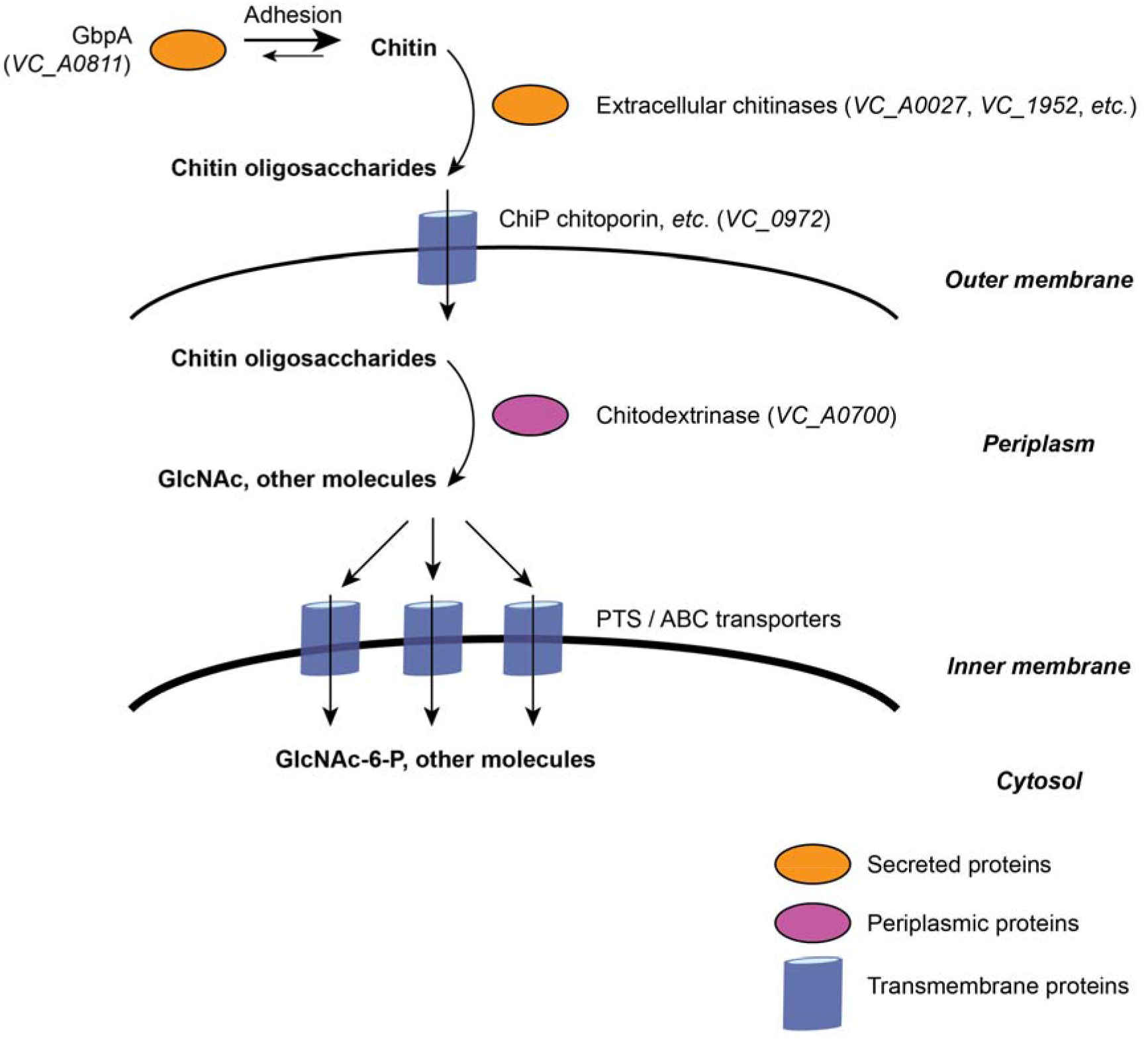
Initial steps in *V. cholerae* chitin uptake and catabolism. Schematic summarising the principal stages in chitin degradation and import by *V. cholerae*. Comprehensive descriptions of this pathway are reported in [19, 35]. The MSHA adhesin has not been included in this diagram.

Previous work used the genomes of 20 diverse *Vibrionaceae* (including seven *V. cholerae)* to determine the presence and absence of genes involved in metabolising chitin across this family of bacteria [19]. However, it is important to note that the chitin degradation pathway of *V. cholerae* has been described using reference strains of the species (particularly in N16961 [23]), and although data exist on how the chitin catabolism pathway varies amongst members of the *Vibrio* genus [19], less is known about how this pathway varies within a single species. This is particularly relevant because emerging evidence suggests that non-7PET lineages of the *V. cholerae* species cause different patterns of disease, even if they harbour some or all of the canonical pathogenicity determinants associated with cholera cases [8]. However, since the chitin metabolic pathway has principally been studied in N16961, a 7PET strain, we know little about the extent to which it varies amongst non-pandemic members of the *V. cholerae* species.

In this study, we focused specifically on genes that encode components of the initial steps of the chitin degradation pathway across the *V. cholerae* species. We focused on these because the functions of many of these genes have been characterised experimentally, and we sought to determine how well the observations in the literature reflect the true distribution of these genes, and their functions, across a diverse species. We generated a pangenome from 195 annotated *V. cholerae* genome sequences, which were chosen to obtain as balanced and unbiased a view of *V. cholerae* as possible (i.e., without focusing solely on epidemic and pandemic lineages). We find that the distribution of these genes is not uniform within *V. cholerae*, and we identify variation amongst the chitinases encoded by diverse *V. cholerae.* We also identify a putatively novel chitinase gene, and present experimental evidence in support of its functional classification.

## Methods

### Strains, plasmids and oligonucleotides

Strains, plasmids, and oligonucleotide primers (Sigma-Aldrich) used for experimental work in this study are listed in Table 1. Bacteria were cultured routinely on LB media supplemented with chloramphenicol (10 μg/ml; LB-Cm) where appropriate.

**Table 1.**
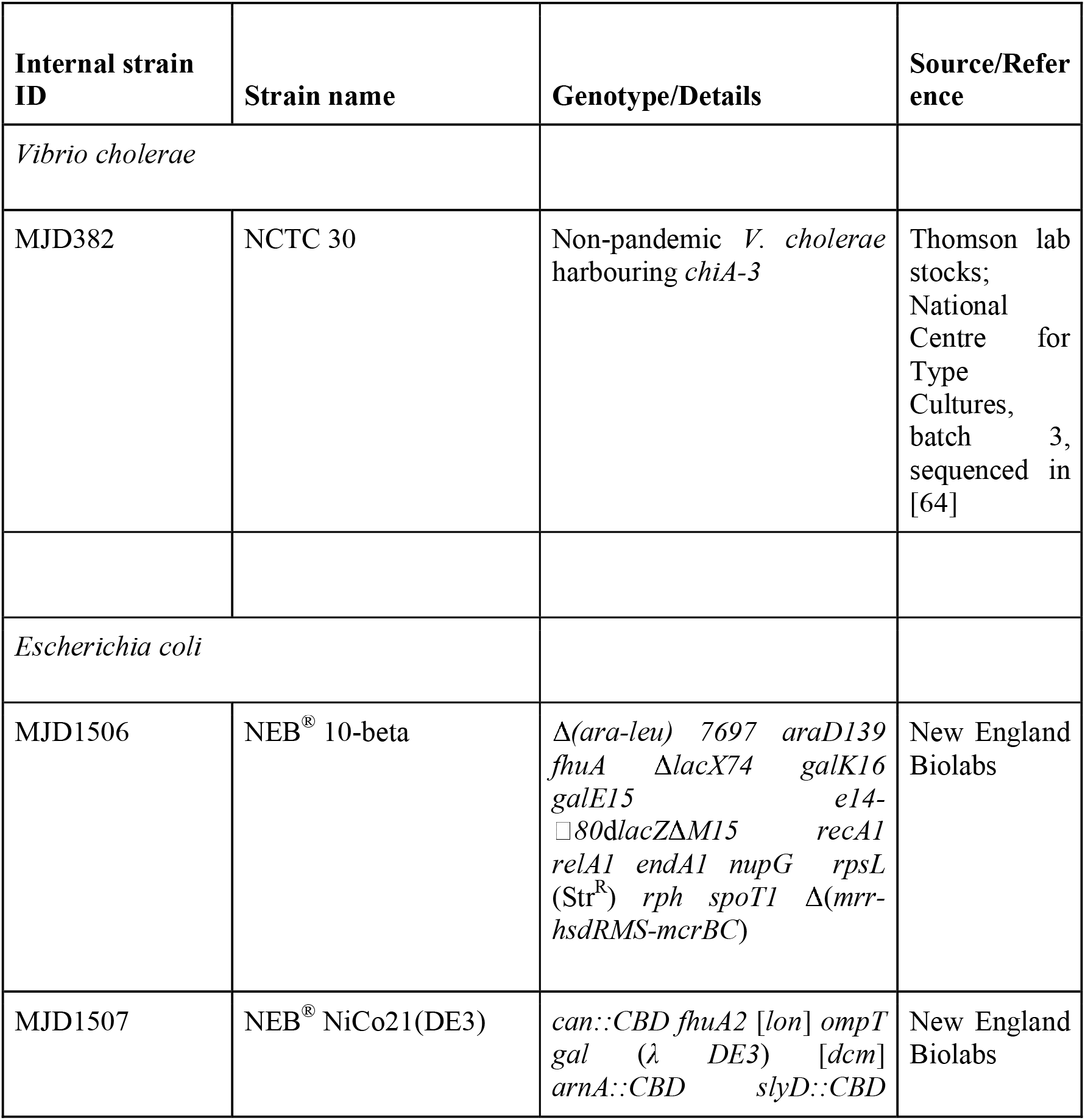
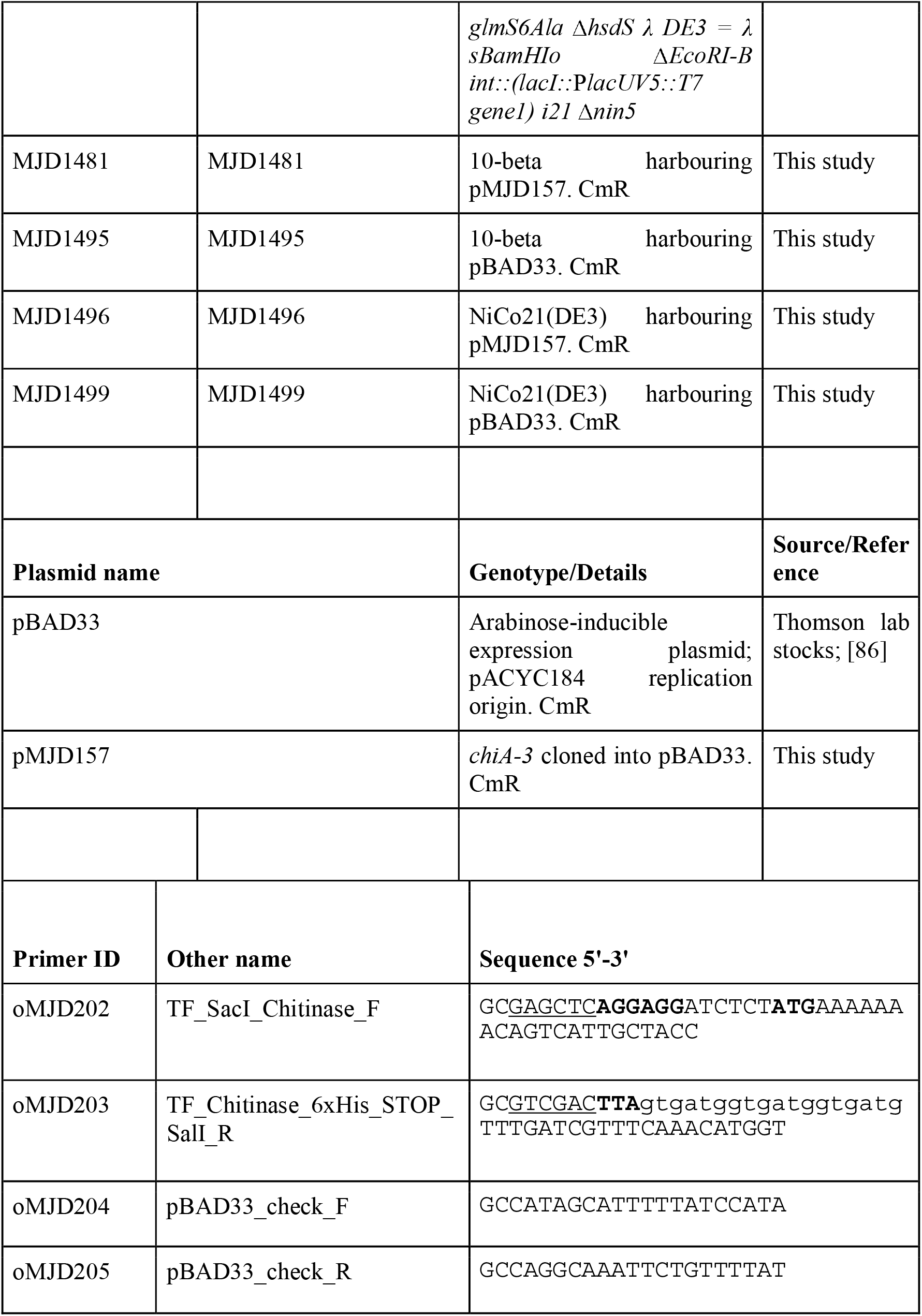
Strains, plasmids, and oligonucleotides used in this study. Restriction enzyme recognition sites are <u>underlined</u>. The primer sequence incorporating a C-terminal 6xHis translational fusion into *chiA-3* is presented in lowercase, and the sequences of ribosome binding sites, start, and STOP codons are in **bold**. CmR = Chloramphenicol resistant. StrR = streptomycin resistant.

### Genome sequences and accession numbers

The 198 genome sequences used to calculate the pangenome described in this manuscript are listed in Supplementary Table 1. Accession numbers for additional chromosome sequences to which the text refers are as follows: *V. harveyi* chromosome 2 (accession # CP009468.1); *V. parahaemolyticus* chromosome 2 (accession # BA000032.2). Accession numbers for the chitinase protein sequences referred to in [19] and used for BLASTp comparisons are listed in Table 2.

**Table 2.**
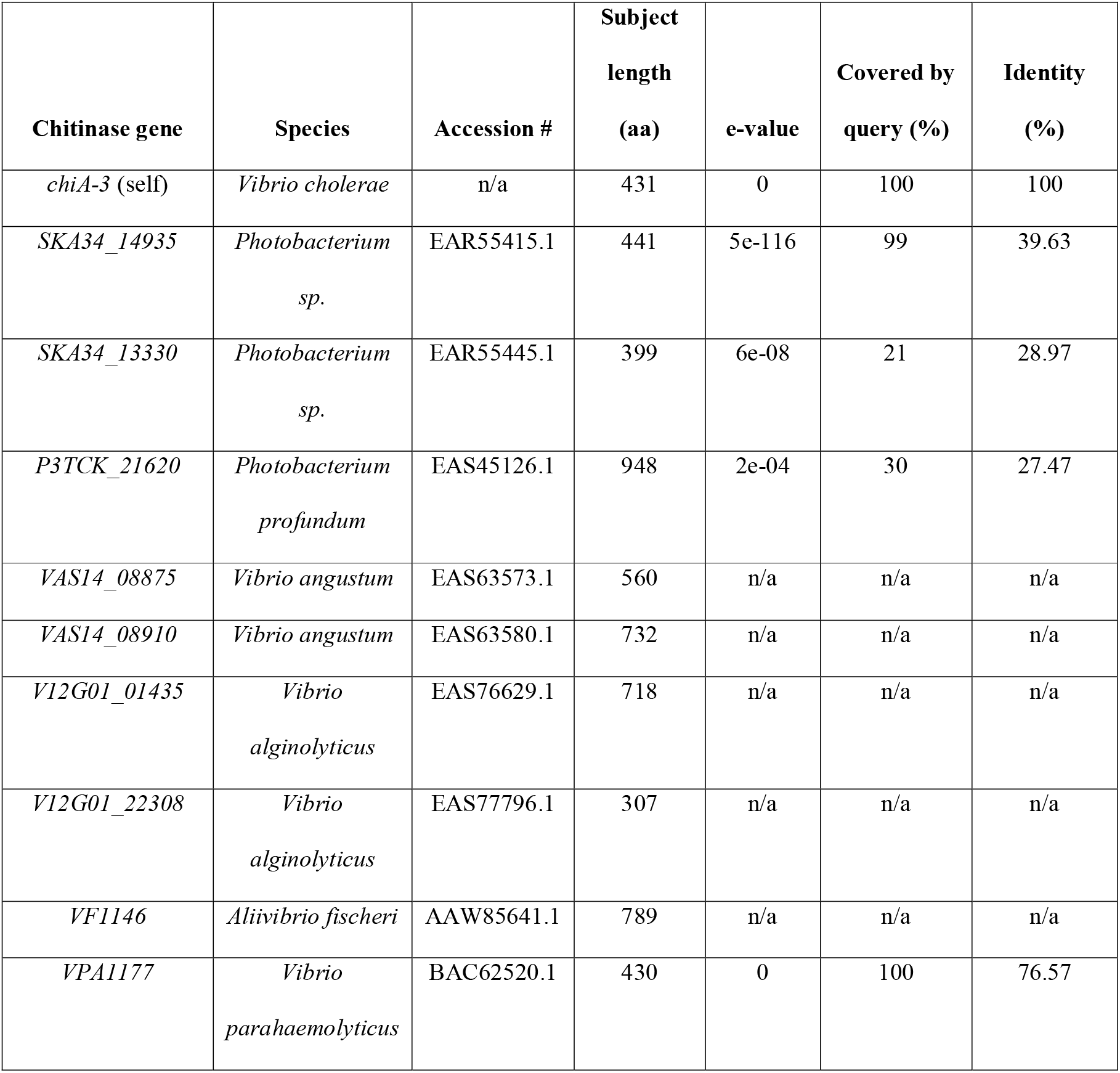
Pairwise BLASTp alignments between *chiA-3* and chitinases from Hunt *et al* [19]. No significant alignment was found between *chiA-3* and *VAS14_08875, VAS14_08910, V12G01_01435, V12G01_22308*, or *VF1146.* A multiple sequence alignment containing each of these protein sequences has been included in the Figshare repository for this study.

### Genome assemblies

*V. cholerae* genome sequences were assembled from short-read data using SPAdes v3.8.2 [49], as part of a high-throughput pipeline [50]. Assemblies were annotated automatically using Prokka v1.5 [51] and a genus-specific reference database [52]. If raw sequencing reads were unavailable for genome sequences, assemblies were downloaded and similarly annotated using the automated Prokka-based pipeline.

### Pangenome and phylogenetic calculations

A pangenome was produced from 198 Prokka-annotated genome assemblies using Roary v3.12.0 [53] (parameters ‘-p 10 -e --mafft -s -cd 97’). A core-gene alignment of 2,520 genes and 1,096,140 nucleotides was produced from this pangenome calculation. The alignment was trimmed using trimAl v1.4.1 [54] and used to produce an alignment of 183,896 SNVs using SNP-sites v2.5.1 [55]. A maximum-likelihood phylogeny was produced using IQ-Tree v 1.6.10 [56] from the SNV-only alignment (options ‘-nt 10 -m GTR+ASC -bb 5000 -alrt 5000’).

### Protein sequence alignments, domain prediction, and comparative genomics

Protein sequences were aligned using BLASTp [57] and were annotated using the InterProScan web server [58]. Comparative genomic figures were generated using BLASTn [57] sequence alignments and visualised using ACT v13 and v18.0.2 [59], and Easyfig v2.2.2 [60].

### Confirmation of gene presence/absence by mapping

Reads were mapped to reference sequences using SMALT v0.7.4 (https://www.sanger.ac.uk/tool/smalt-0/) and the method described by Harris et al. [61], as part of automated analysis pipelines run by Wellcome Sanger Institute Pathogen Informatics. All of the software developed by Pathogen Informatics is freely available for download from GitHub under an open source license, GNU GPL 3 (https://github.com/sanger-pathogens/vr-codebase). Ordered BAM files were visualised against reference sequences using Artemis v16 and v18.0.2, which incorporates BamView [62, 63].

### Molecular cloning

Plasmid DNA was extracted from *E. coli* using the QIAprep Spin Miniprep kit (Qiagen, #27104). Genomic DNA (gDNA) was extracted from NCTC 30 as described previously [64]. Cloning intermediates were purified using the QIAquick PCR Purification kit (Qiagen, #28104).

gDNA from NCTC 30 was used as a template from which to amplify *chiA-3* using primers oMJD202 and oMJD203, high-fidelity Phusion Hot Start Flex polymerase (NEB #M0535S) using the supplied GC buffer, DMSO (3% v/v final conc.) and dNTPs (Thermo Scientific, #R0191). Twenty-nine PCR cycles were performed using the manufacturer’s protocol (annealing temperature: 55 °C, extension time: 2 min). The amplicon was purified and digested using 30 units of SacI-HF and SalI-HF (NEB, #R3156S and R3138S respectively) at 37 °C for 45 min. pBAD33 was similarly treated with SacI-HF and SalI-HF, and after 15 min incubation at 37 °C, the plasmid digestion was supplemented with 1.5 units of recombinant shrimp alkaline phosphatase (rSAP; NEB #M0371S). Digested insert and vector were purified and ligated together at room temperature for 30 min using T4 DNA ligase (NEB, #M0202S) in approximately a 3:1 molar ratio. Chemically competent 10-beta *E. coli* (NEB, #C3019I) were transformed with ligated DNA according to the manufacturer’s instructions, and transformants were selected for on LB agar supplemented with chloramphenicol (10 μg/ml).

Chloramphenicol-resistant colonies were resuspended in 30 μl PBS. A screen for clones containing an insert into pBAD33 was carried out using 1 μl of this suspension as a PCR template using primers oMJD204 and oMJD205 and OneTaq Quickload 2X Master Mix (NEB, #M0486S), according to the manufacturer’s instructions (annealing temperature 45 °C, extension time 3 min). Plasmids were extracted from overnight cultures of clones from which PCR produced an amplicon of the expected size (1,548 bp). The presence of an insertion into pBAD33 was verified by digesting purified plasmid DNA with SacI-HF and SalI-HF as described above. Plasmids were then sequence-confirmed by amplicon sequencing (GATC/Eurofins) in both directions across the pBAD33 multiple cloning site using primers oMJD204 and oMJD205. Sequence-verified plasmids were transformed into chemically competent NiCo21(DE3) cells (NEB, #C2529H) following the manufacturer’s instructions, and these transformants were used for protein expression purposes.

### Protein expression and immunoblotting

Single colonies of NiCo21(DE3) harbouring pMJD157 and pBAD33 (empty vector) were inoculated into 3 ml LB-Cm and cultured at 37 °C with shaking (200 rpm) for eight hours. These were then diluted 1:100 into baffled flasks containing 25 ml LB-Cm supplemented with either □-(+)-glucose (BDH, #101176K) or □-(+)-arabinose (Sigma-Aldrich, #A3256), both at 0.4% w/v final concentration. These cultures were grown for 18 hours at 23 °C with shaking (200 rpm). Cells were collected by centrifugation (3,900 x *g*, 5 min) and the supernatant was filter-sterilised (0.22 μm) and stored at −20 °C. Cell pellets were lysed in 3 ml BugBuster HT (Millipore, #70922-4) for 30 min at room temperature on a rotator. Debris was collected by centrifugation (3,900 x *g*, 5 min) and discarded. Lysates were stored at −20 °C.

Sixty microlitres of filtered supernatants and lysates was mixed 1: 1 with 2X tris-glycine-SDS sample buffer (Invitrogen, #LC2676), boiled at 100 °C for 5 min, and 30 μl of each sample was used to load duplicate NuPAGE 4-12% Bis-Tris acrylamide gels (Invitrogen, #NP0321) which were electrophoresed simultaneously, in the same gel tank. Stained and unstained protein ladders (NEB; #P7719S and #P7717S; Invitrogen, #LC5925) were used for size estimation where appropriate. One gel of the pair was stained with InstantBlue (Expedon, #ISB1L) according to the manufacturer’s instructions prior to imaging; the other was used for Western immunoblotting.

For Western blotting, electrophoresed proteins were transferred from an acrylamide gel onto a nitrocellulose membrane using the iBlot 2 dry blotting system and transfer stack (ThermoFisher, #IB21001 and #IB23001). After transfer, the membrane was blocked for three hours in 5% w/v Marvel milk powder dissolved in PBS-Tween 20 (Marvel-PBS-T) at 4 °C, with rocking. An antibody recognising the 6xHis epitope and directly conjugated to horseradish peroxidase (Abcam, #ab1187) was diluted in Marvel-PBS-T to the manufacturer’s instructions and used to probe the membrane for 30 min at 4 °C, with rocking. The membrane was then washed in PBS-T for 15 min three times, and then incubated with Clarity Western ECL substrate (Bio-Rad, #170-5060) for 5 min. Luminescence signal was allowed to decay overnight, and the blot was then imaged with Amersham Hyperfilm ECL film (GE, #28906836). Coloured protein size standards were marked manually on the developed film.

### Chitinase assay

Chitinase activity was assayed using fluorogenic substrates (Sigma-Aldrich, #CS1030). The kit was used according to the manufacturer’s instructions, with the following modifications: Ten microlitres of cell lysate or supernatant was used per assay well. Five microlitres of the supplied chitinase control enzyme was used per positive control reaction, rather than a 1:200 dilution of the control enzyme, to ensure that fluorescence was detectable. Assays were carried out in black Nunc flat-bottomed microtitre plates (Sigma-Aldrich, #P8741), and technical triplicates were included for each sample. Once mixed, reaction plates were incubated for 30 min (37 °C, static) before the addition of stop solution. Fluorescence was measured using a FLUOstar^®^ Omega plate reader (BMG LabTech), set to excitation and emission wavelengths of 360 and 450 nm, respectively. A 1% gain was applied to the fluorescence measured by the reader. Blank fluorescence was subtracted from each sample reading prior to analysis.

### Statistics, data visualisation, and figure generation

Figures were produced using R v3.5.1 [65], ggpubr v0.2.3 (https://github.com/kassambara/ggpubr), ggplot2 v3.2.1 [66], ggforce v0.3.1.9000 (https://github.com/thomasp85/ggforce), and the Phandango web server [67]. Statistical tests were performed using R v3.5.1 [65]. Where required, figures were modified manually using InkScape v0.92.4 and Adobe Illustrator CC v23.1.1.

## Results

### Distribution of chitinase genes amongst *V. cholerae*

The key components of *V. cholerae* chitin catabolism summarised in Figure 1 have been previously described [23, 44, 68]. The presence and absence of orthologues of each of the principal chitin-binding proteins and extracellular chitinases [44] known to be encoded by the *V. cholerae* 7PET reference strain N16961 (based on their N16961 locus identifiers) were identified in a pangenome calculated from 195 *V. cholerae* genomes, plus three *Vibrio* spp. genomes used as an outgroup (Supplementary Table 1). Genes that were annotated as encoding putative chitinases, as well as those genes known to be present in N16961, were identified in the pangenome (Supplementary Tables 2, 3). A *V. cholerae* phylogenetic tree was calculated using an SNV-only alignment of 2,520 core genes taken from the pangenome, and the distribution of these chitinase genes across the phylogeny is presented in Figure 2.

**Figure 2.**
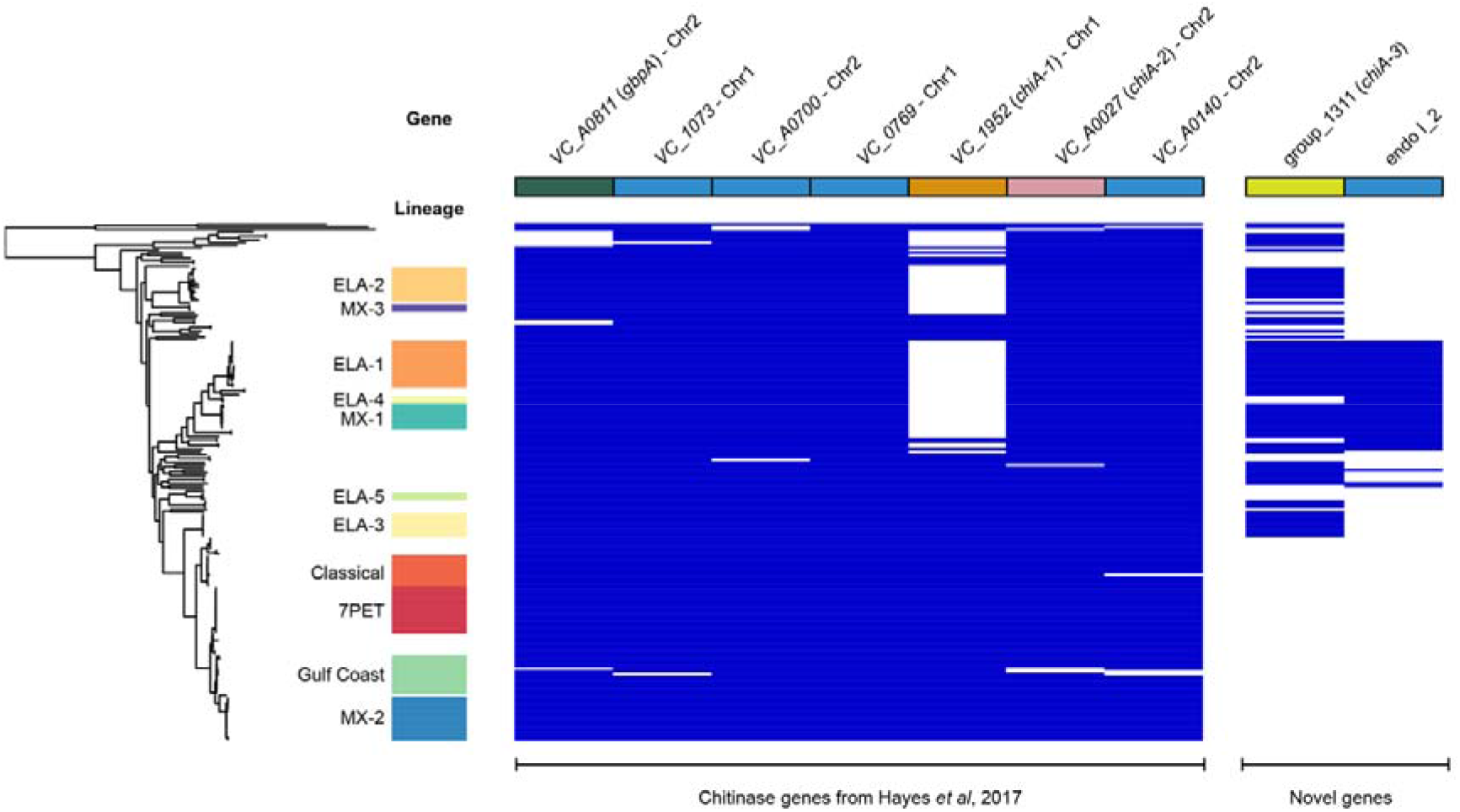
Distribution of chitinase genes amongst diverse *V. cholerae*. Visualisation of the presence and absence of genes encoding key *V. cholerae* chitinase enzymes and chitin adhesion factors (Figure 1). The seven genes encoding putative *V. cholerae* endochitinases described by [44] are listed, as well as two additional putative chitinases identified in this analysis. Figure generated using Phandango [67]. Isolate assignments to *V. cholerae* lineages were taken verbatim from [64, 87], and are named after [8]. Chromosomal location assigned to genes present in N16961. Colour coding of *chiA-1, chiA-2*, and *chiA-3* is consistent among figures in the manuscript.

### *gbpA* is not universally present amongst diverse *V. cholerae*

The first, and most striking, observation made from these data was that *gbpA (VCA_0811)* did not appear to be ubiquitous amongst all of the *V. cholerae* genomes included in this study. We found that *gbpA* was present in only 189 of 195 *V. cholerae* genomes (96.9%; Figure 2, Supplementary Table 2). We manually inspected the genome assembly for each isolate which lacked *gbpA*, to guard against this being an artefact of the computational approach taken (Figure 3).

**Figure 3.**
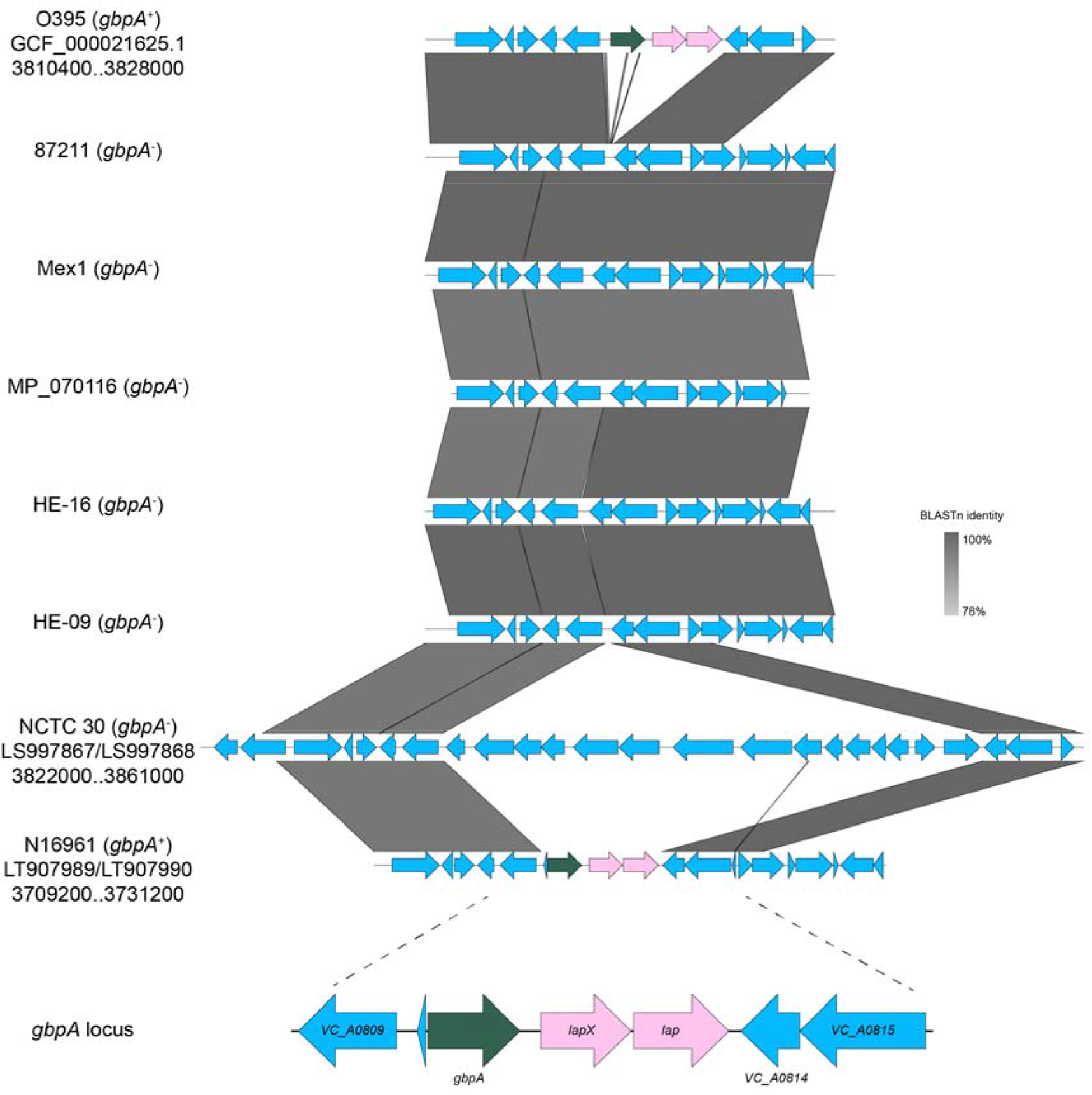
Confirming the absence of *gbpA* and adjacent genes from assemblies. Heterogeneity at the genomic locus encompassing *VC_A0811* was observed in isolates lacking gbpA. All assemblies lacked *VC_A0811-VC_A0813*, and these genes were replaced with sequence containing at least 15 genes in NCTC 30. Read mapping data confirming the absence of *VC_A0811-VC_A0813* from NCTC 30 are presented in Supplementary Figure 1. Accession numbers and assembly co-ordinates are reported for reference and closed genome sequences. Figure generated using Easyfig [60] and BLASTn comparisons [57]. *VC_A0811-VC_A0813* are highlighted in N16961 and O395 (both *gbpA^+^)* for ease of illustration.

Three genomic arrangements were observed at this locus – the presence of an intact *VC_A0811* locus as found in *gbpA+* genomes, a deletion of *gbpA* and two adjacent genes (*VC_A0811*-*VC_A0813*), and the replacement of these three genes with additional sequence in the genome of NCTC 30 (Figure 3). In order to ensure that the *VC_A0811*-*VC_A0813* genes were not present at a different position in the NCTC 30 genome, we mapped the Illumina short-reads for this isolate to the N16961 reference sequence and inspected the mapping coverage across this region. This confirmed that the absence of the genes *VC_A0811-VC_A0813* from NCTC 30 was not a result of a mis-assembly (Supplementary Figure 1). The two genes adjacent to *gbpA, VC_A0812* and *VC_A0813*, encode LapX and Lap, respectively. Both genes are putatively regulated by the HapR master quorum-sensing regulator, and encode proteins that were detected in an *hapA* mutant [69]. Both Lap and LapX were found to be putative components of the Type 2 secretome in N16961 [70], and *lap* has been used as a polymorphic locus in multilocus enzyme electrophoresis MLEE schemes for classifying *V. cholerae* [71, 72]. We were unable to find published evidence linking these genes to GbpA activity or chitin adhesion more generally, though we note that *lap* and *lapX* are oriented in the same direction as *gbpA*, and we cannot exclude the possibility that these three genes are co-regulated or co-transcribed.

### *chiA-2*, but not *chiA-1*, is ubiquitous amongst diverse *V. cholerae*

In contrast to *gbpA*, we found that *VC_A0027* (encoding ChiA-2) was near-ubiquitous, being detected in 192/195 *V. cholerae* (Figure 2; Supplementary Table 2). Manual inspection of the assemblies for those three isolates which appeared to lack the gene confirmed that the majority of this gene was in fact present; assembly and resultant annotation errors were likely to be responsible for this result (data not shown). This suggests that *VC_A0027* is core to *V. cholerae*, which is consistent with this being the most highly-expressed chitinase enzyme in the species, and with the observation that deletion of this gene alone causes a significant growth defect on minimal media containing chitin as a sole carbon source [44].

However, although *VC_1952* (ChiA-1) was present in all pandemic isolates (defined as those isolates which were members of the 7PET and Classical lineages), it was not ubiquitous across the species, and was only identified in 61.2% of the non-pandemic *V. cholerae* in this dataset (101/165; Figure 2; Supplementary Table 2). This observation was surprising, because both ChiA-1 and ChiA-2 have been shown to be necessary for *V. cholerae* N16961 to grow in media supplemented with colloidal chitin [23]. Keymer and colleagues previously observed, using microarray approaches, that some diverse environmental isolates of *V. cholerae* varied in terms of their *VC_1952* genotype [73]. We propose that our data recapitulate this observation, albeit *in silico*. We manually examined the region surrounding the *VC_1952* locus in a subset of the genome assemblies for isolates lacking this gene, and found both that the gene was absent in its entirety, and that this did not appear to affect the genes adjacent to *chiA-1* (Figure 4; Supplementary Figure 2). Moreover, the distribution of putative chitinases (Figure 2) suggested that isolates lacking ChiA-1 may encode additional chitinases. Since ChiA-1 is known to have a functional role in *V. cholerae* chitin metabolism, this led us to speculate that these additional putative chitinases, if functional, might be able to provide chitinase activity in the absence of ChiA-1.

**Figure 4.**
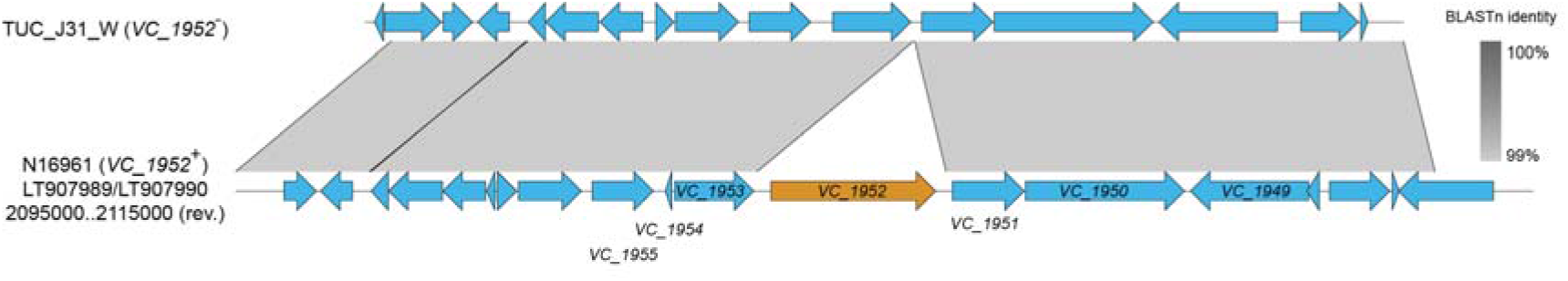
Genes adjacent to the *VC_1952* locus are intact in genomes lacking *chiA-1*. An example is presented in which the genes flanking *VC_1952* remain intact in the absence of *VC_1952* itself, contrasting with the observation made at the *gbpA* locus (Figure 3). A larger number of diverse genomes are similarly analysed in Supplementary Figure 3. Figure generated using Easyfig [60] and BLASTn comparisons [57].

### Identification and characterisation of *chiA-3*

Eleven gene clusters in the pangenome included genes with the annotation “chitinase” or “putative chitinase” (Supplementary Table 3). Five of these were found only in one genome, of which four were found only in the non-*V. cholerae* outgroup. Of the remaining six genes, four are known to be present in N16961 (Supplementary Tables 2 and 3). On further examination, the products of one of the two gene clusters, ‘endo I_2’, were not predicted *in silico* to contain a chitinase domain, although a putative chitin-binding domain was identified (Supplementary Figure 3; Supplementary Table 3).

The second gene identified was predicted to encode a protein containing a chitinase domain (Figure 5a). The molecular weight (47.69 kDa) and domain composition of the protein were distinct from those of *chiA-2* and *chiA-1* (Figure 5a), as was the genomic context and location of the gene, which was inserted between *VC_A0620* and *VC_A0621* on chromosome 2 (Figure 5b). This gene was therefore referred to as *chiA-3*, to differentiate it from the two previously-described genes. *chiA-3* was identified in 87 genomes, and was absent from all of the genomes belonging to both pandemic *V. cholerae* lineages included in this study. Additionally, 57 of the 67 isolates which lacked *chiA-1* harboured *chiA-3* (85.0%).

**Figure 5.**
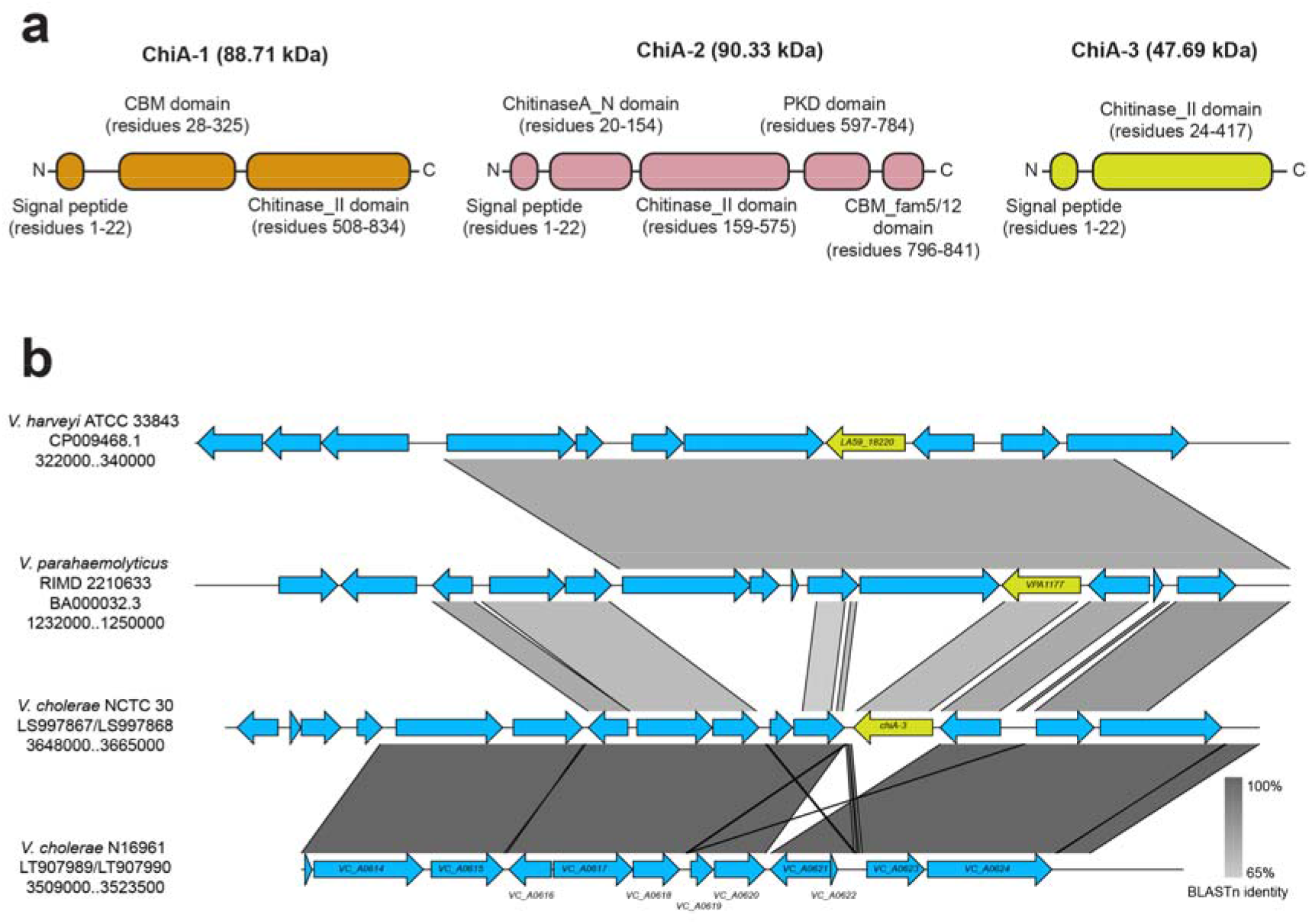
ChiA-3 is a protein distinct from ChiA-1 and ChiA-2. (a): Cartoons of the protein domains predicted to be present in each of ChiA-1, ChiA-2, and ChiA-3 are presented. Predicted molecular weights are indicated. Proteins are not to scale. (b): *chiA-3* is integrated between *VC_A0620* and *VC_A0621* in the smaller *V. cholerae* chromosome. This genomic position is conserved in *Vibrio* spp. which harbour *chiA-3* orthologues [74, 77]. Figure generated using Easyfig [60] and BLASTn comparisons [57].

In order to determine whether *chiA-3* had been identified previously in other *Vibrio* species, the gene was compared to the nine genes listed by Hunt *et al* as chitinases found in non-cholera Vibrios [19] (Table 2). The most similar protein to ChiA-3 (76.57% aa identity) was that encoded by *VPA1177 (chiA*, accession # BAC62520.1), found in *V. parahaemolyticus* strain RIMD 2210633 (Table 2) [74]. *VPA1177* encodes a 430 aa protein (47.98 kDa) which previous genetic analyses have shown to make a minimal contribution to the ability of *V. parahaemolyticus* to degrade chitin - ChiA-2 (encoded by *VPA0055*, accession # BAC61398.1) is the major protein responsible for this phenotype in *V. parahaemolyticus* [75]. Transcription of *VPA1177* has been shown to be significantly reduced in the presence of chitin [75], however, the VPA1177 protein has been shown to be expressed by *V. parahaemolyticus*, albeit at very low levels in culture supernatants [35].

A previous report had also identified a functional secreted chitinase from *Vibrio harveyi* of a similar molecular weight (47 kDa) to both VPA1177 and ChiA-3 [76]. The *V. harveyi* ATCC 33843 genome [77] contains a gene encoding a putative chitinase (BLASTp: 100% query coverage, 77.73% amino acid identity to ChiA-3; predicted molecular weight 48.0 kDa) in a similar genomic context on chromosome 2 to that of *chiA-3* in NCTC 30 (Figure 5b). This is distinct from the location of the functionally-characterised *chiA* gene *(LA59_20935)* which encodes an 850 aa ChiA chitinase precursor (accession # Q9AMP1 [36, 78, 79]), and from other functionally-characterised *V. harveyi* β-*N*-acetylglucosaminidases [37]. This *V. harveyi* protein is also 90.9% identical to VPA1177. As well as their high amino acid identity, each of these proteins were predicted to contain similar domain compositions and configurations across the three species (Supplementary Figure 4). It is reasonable to infer that these enzymes are orthologues of ChiA-3.

Since *VPA1177* has been shown to be transcribed [75] and to produce a translated protein in *V. parahaemolyticus* [35], we sought to determine whether the product of *chiA-3* from *V. cholerae* had chitinase activity. We amplified the gene from the genome of NCTC 30, a non-pandemic lineage *V. cholerae*, and cloned it directionally into pBAD33 such that expression of the gene was governed by the arabinose-inducible P_BAD_ promoter and the translated product linked to a C-terminal 6xHis tag, similar to previous reports [39, 44] (denoted pMJD157, Figure 6a).

**Figure 6.**
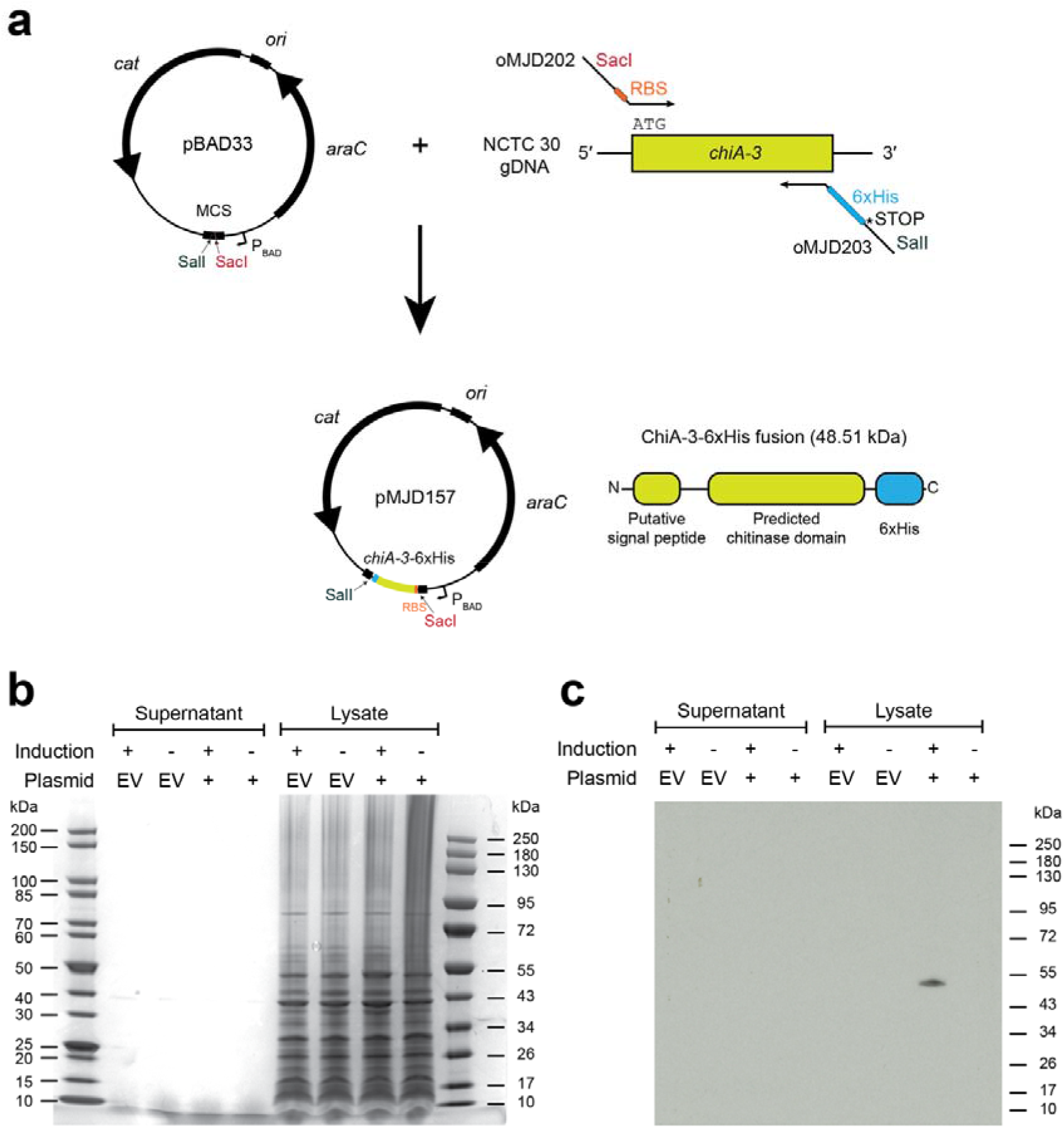
Molecular cloning of *chiA-3* and expression of ChiA-3-6xHis. (a): Schematic of cloning strategy used to amplify and insert *chiA-3* directionally into the pBAD33 multiple cloning site (MCS), under the arabinose-inducible P_BAD_ promoter, and to incorporate a C-terminal 6xHis tag as a translational fusion. A linker sequence was not incorporated between the C-terminus of ChiA-3 and the 6xHis tag. Figures are not to scale. (b): InstantBlue-stained acrylamide gel of proteins present in supernatants and cell pellet lysates from cultures grown at 23 °C supplemented with arabinose (induction +) or glucose (induction -). No induced bands were easily discerned. (c): Western immunoblot produced from an identically-loaded acrylamide gel to that presented in (b), run in parallel with the gel in (b), and probed with an α-6xHis antibody (see Methods). A band corresponding to the expected molecular weight of ChiA-3-6xHis (48.51 kDa) was detected in the cell pellet lysate of *E. coli* harbouring pMJD157 only (plasmid +). This size is consistent with the retention of the fusion protein without the cleavage of the putative signal sequence. Protein ladders: NEB #P7719S and #P7717S. EV = empty vector (pBAD33).

*E. coli* harbouring pMJD157 produced a His-tagged protein of the expected molecular weight that was retained in the cell pellet when cultured with arabinose at 23 °C (Figure 6b). We used a commercial fluorogenic assay for chitinase activity which relies on the hydrolysis of 4-methylumbelliferyl (4-MU) chitin analogues to detect chitinase activity. A similar assay has been used previously to assay chitinase activity in Vibrios [76]. We found that samples from *E. coli* cultures expressing 6xHis-tagged ChiA-3 demonstrated statistically significant activity on 4-MU-linked substrates (Figure 7). These data were consistent with the His-tagged protein detected in Figure 6c (ChiA-3-6xHis) having endochitinase and chitobiosidase activities, but lacking β-*N*-acetylglucosaminidase activity.

**Figure 7.**
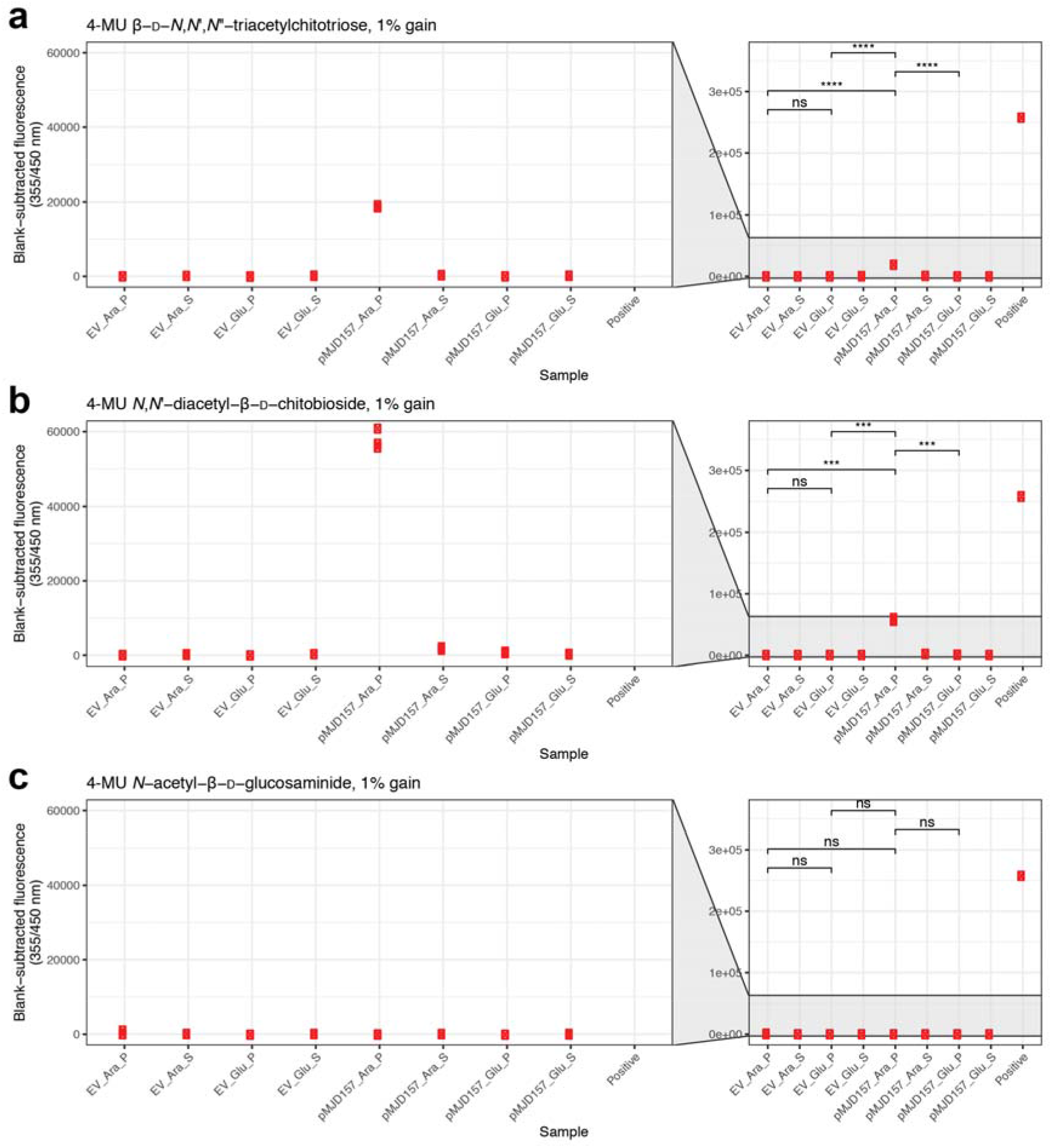
ChiA-3-6xHis displays chitobiosidase and endochitinase activities, but not β-*N*-acetylglucosaminidase activity. Lysates and supernatants included in Figures 6b and 6c were assayed for chitinase enzyme activity using a fluorometric chitinase assay kit (see Methods for details). Lysed cells from *E. coli* cultures harbouring pMJD157 and cultured in the presence of arabinose were the only samples which produced detectable and statistically significant signals on triacetylchitotriose and chitobiose substrates (a, b). No signal was detected in the presence of glucosaminide substrate (c). All plots are scaled equivalently. P = pellet; S = supernatant; EV = empty vector (pBAD33). Parametric t-tests performed where indicated: ns = not significant; *** = p<0.001; **** = p<0.0001.

## Discussion

In this study, we present three major observations - firstly, that *gbpA* is not ubiquitous amongst *V. cholerae.* Second, we show that there is additional variability in the chitinase genes harboured by diverse *V. cholerae*, which show phylogenetic signals in their distribution. Third, we present functional evidence that one of these putatively-novel genes encodes a protein with chitinase activity.

The fact that *gbpA* is not present in all *V. cholerae* is important, given that *gbpA* had previously been suggested to be a candidate diagnostic gene for the detection of *V. cholerae* [80, 81]. This was based both on the high level of conservation of *gbpA* amongst tested *V. cholerae*, and on the number of differences between *gbpA* in *V. cholerae* and alleles found in other *Vibrio* species [80]. In addition to our results, others have noted that *gbpA* can be found in non-cholera Vibrios and in non-pathogenic *V. cholerae*, suggesting that this makes *gbpA* an unreliable marker for quantitative study of *V. cholerae* [82].

The biological consequences of the absence of *gbpA* from these bacteria is interesting to consider. As discussed previously, GbpA is an important factor in both environmental and pathogenic colonisation. The fact that *gbpA* is absent from non-pandemic *V. cholerae* that appear to be basal to the rest of the species (Figure 2) suggests that its role in pathogenicity may be more complex than previously thought. It might be that acquisition of *gbpA* by *V. cholerae* was an important step in its evolution as a human pathogen. Conversely, since several of the isolates in the lineage lacking *gbpA* are of clinical as well as environmental origin [8, 64, 83–85], including some which were isolated from cases of acute or ‘choleraic’ diarrhoea [64, 83, 84], it might be that *gbpA* may not be essential for pathogenic colonisation. It remains to be seen whether the natural absence of *gbpA* affects the ability of such *V. cholerae* to colonise both the intestinal mucosa and chitinous surfaces. The roles played by other adhesins in these diverse *V. cholerae*, such as MSHA, should also be considered in the future.

Although ChiA-3 orthologues have been examined in other Vibrios, we believe that this is the first report of this gene in *V. cholerae*, and the first report that the *V. cholerae chiA-3* gene encodes a functional chitinase. The fact that *chiA-3* was found only in non-pandemic *V. cholerae* is also intriguing. It is not yet known whether non-pandemic *V. cholerae* harbouring *chiA-3* can respire chitin as effectively, or more effectively, than N16961 or other laboratory strains. However, it could be speculated that ChiA-3 might be more suited to environmental survival than ChiA-1 (e.g., lower temperatures, higher salinity than the human intestine). Although such investigations were outside the scope of this current study, quantifying the relative activities of ChiA-3 and ChiA-1, and determining genetically whether *chiA-3* can complement the loss of *chiA-1* from *V. cholerae*, or if isolates that harbour both *chiA-1* and *chiA-3* (Figure 2; Supplementary Table 1) have an enhanced chitin degradation phenotype, is the subject of future research.

There are fundamental differences between *V. cholerae* from pandemic and non-pandemic lineages, both in terms of their ability to cause cholera epidemics, and their basic biology. We still do not fully understand these differences, but in order to do so, we must study *V. cholerae* pathogenicity in conjunction with more fundamental biological processes. It is currently unclear whether variation in the complements of chitinases and chitin-binding proteins encoded by *V. cholerae* have physiological consequences for different lineages of the species. However, given the importance of these genes to pathogenicity [29, 39], environmental lifestyles [23], and natural competence [28], it is plausible that these differences reflect differences in the ecological niches occupied by different lineages of the species. Research in this area may provide further insights into the genetic and biochemical differences between *V. cholerae* lineages that cause dramatically different patterns of disease worldwide. Collectively, these findings underline the fact that, as we continue to study diverse *V. cholerae*, our understanding of the nuance and specifics of this species will improve and be refined.

## Author statements

### Author contributions

N.R.T. supervised the work. T.G.F. performed genomic analysis with assistance from M.J.D. and G.A.B.. M.J.D. carried out experimental work. T.G.F. and M.J.D. wrote the manuscript, with major contributions from N.R.T.. All authors interpreted the results, contributed to the editing of the manuscript, and read and approved the final version of the manuscript.

## Acknowledgements

We thank Rita Monson for helpful discussions and comments throughout the project. We thank Sally Kay for logistical support, Charlotte Tolley for the gift of reagents, and the Wellcome Sanger Institute (WSI) Pathogen Informatics team for help with data management. We also thank the WSI Parasites and Microbes Administration team, particularly Kate Auger and Joseph Woolfolk, for operational and administrative assistance in the course of this work.

## Abbreviations

4-MU: 4-methylumbelliferyl
ABC transporter: ATP-binding cassette transporters
CmR: chloramphenicol resistant
DMSO: dimethyl sulfoxide
dNTP: deoxyribonucleoside triphosphate
GlcNAc: *N*-acetyl-β-d-glucosamine
MSHA: mannose-sensitive haemagglutinin
SNV: single nucleotide variant
StrR: streptomycin resistant
PTS transporter: phosphotransferase transporter.

