## Supplementary Material for "*gbpA* and *chiA* genes are not uniformly distributed amongst diverse *Vibrio cholerae*"

These supplementary materials include:

**Supplementary Table 1** (attached .xls)

**Supplementary Tables 2-3**

**Supplementary Figures 1-4**

**Supplementary References**

Additional materials that support this study are available in Figshare:

https://dx.doi.org/10.6084/m9.figshare.13169189

(Note for peer-review: Figshare DOI is inactive but will be activated upon publication, please use temporary URL https://figshare.com/s/7795a2d80c13f694f8fa for review)

**Supplementary Table 2.** Summary of seven previously-published putative and validated chitinase genes from N16961 [1] annotated in the *V. cholerae* pangenome. Order corresponds to that of Figure 2. Gene cluster IDs correspond to the gene presence/absence matrix available in the Figshare repository supporting this study. The dataset contains a total of 198 genomes, of which three are not *V. cholerae*.

| **Gene cluster ID** | **N16961 locus ID** | **# isolates containing gene cluster (198 total)** | **Details** |
|---|---|---|---|
| gbpA | *VC_A0811* | 189 | Encodes GbpA |
| endo I_1 | *VC_1073* | 196 | Putative chitinase |
| group_316 | *VC_A0700* | 195 | Encodes chitodextrinase |
| endo I_3 | *VC_0769* | 198 | Putative chitinase |
| chiD | *VC_1952* | 131 | Encodes ChiA-1 |
| chiA | *VC_A0027* | 194 | Encodes ChiA-2 |
| chiA_2 | *VC_A0140* | 194 | Putative chitinase |

**Supplementary Table 3.** All genes annotated in the *V. cholerae* pangenome as 'chitinase' or 'putative chitinase'. Gene cluster IDs correspond to the gene presence/absence matrix available in the Figshare repository supporting this study. The dataset contains a total of 198 genomes, of which three are not *V. cholerae*.

| **Gene cluster ID** | **Annotation** | **# isolates containing gene cluster (198 total)** | **N16961 locus ID** |
|---|---|---|---|
| endo I_3 | putative chitinase | 198 | *VC_0769* |
| endo I_1 | putative chitinase | 196 | *VC_1073* |
| chiA | chitinase | 194 | *VC_A0027* (*chiA-2*) |
| chiD | chitinase | 131 | *VC_1952* (*chiA-1*) |
| group_1311 | chitinase | 87 | n/a (*chiA-3*) |
| endo I_2 | chitinase | 45 | n/a (no chitinase domain) |
| group_14144 | chitinase | 1 | n/a |
| VCJ_003148 | chitinase | 1 | n/a |
| VCJ_001093 | chitinase | 1 | n/a |
| VOA_002115 | chitinase | 1 | n/a |
| VOA_001161 | chitinase | 1 | n/a |

### Supplementary Figures

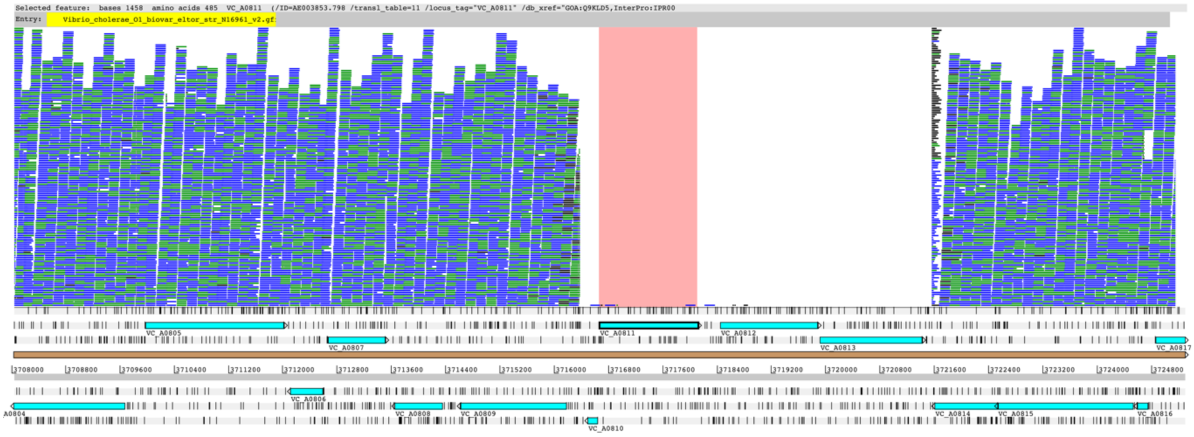


**Supplementary Figure 1. Confirming the absence of *VC_A0811-VC_A0813* from NCTC 30 by mapping.** Illumina short-reads from NCTC 30 were mapped to the N16961 reference sequence and visualised using Artemis and BamView [2, 3]; *gbpA* is highlighted. The drop in coverage over these genes indicates that these are absent from the sequenced NCTC 30 genome.

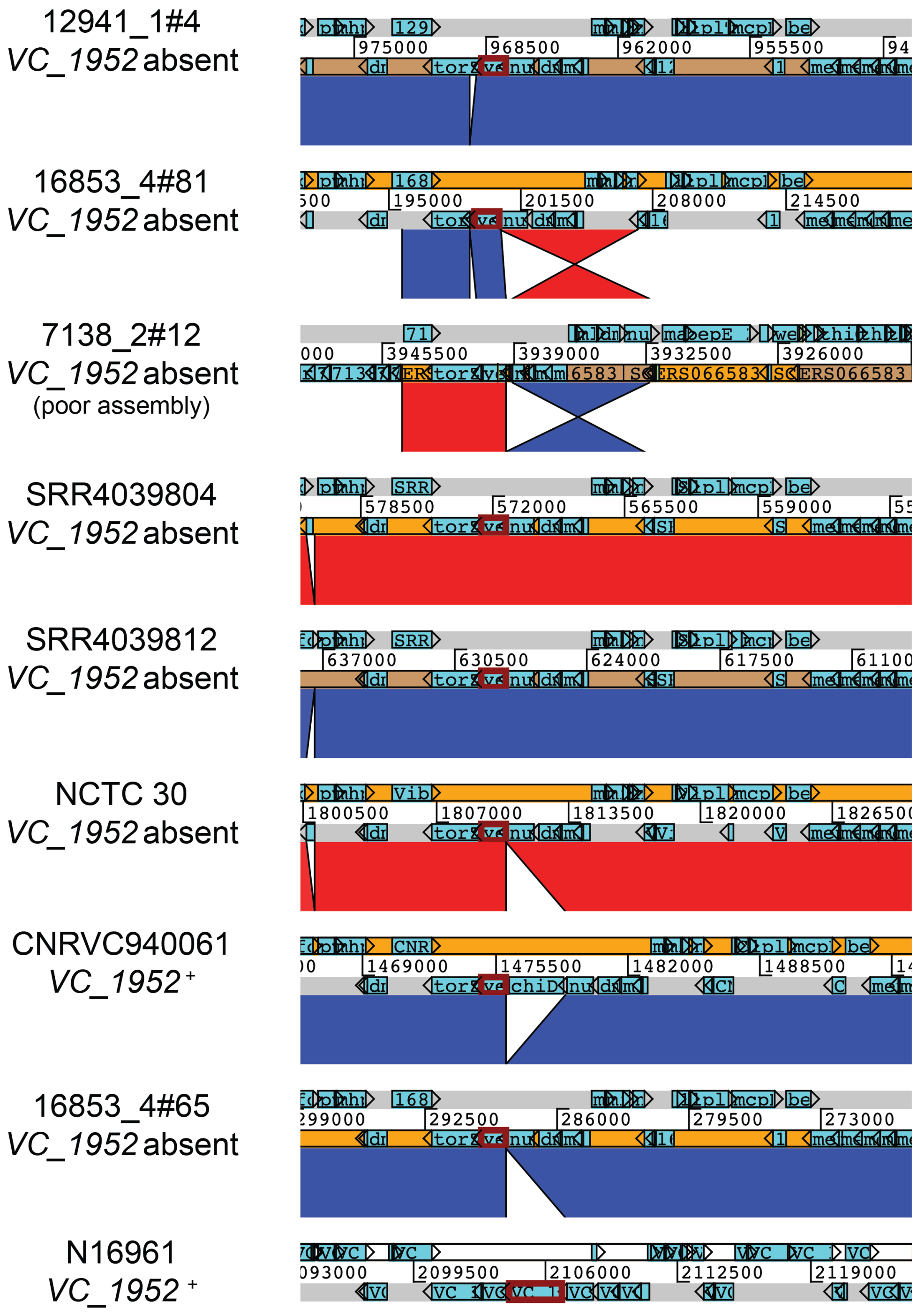


**Supplementary Figure 2. Loss of *VC_1952* in multiple diverse *V. cholerae*.** ACT comparisons of assemblies for a set of seven *V. cholerae* lacking *chiA-1* and two which harbour *chiA-1*, aligned using BLASTn [4]. In all cases, the absence of *chiA-1* (*VC_1952*) does not interfere with the genes surrounding this locus. The *chiA-1* gene has been highlighted in the N16961 reference sequence (bottom of figure), and syntenic contigs have been aligned.

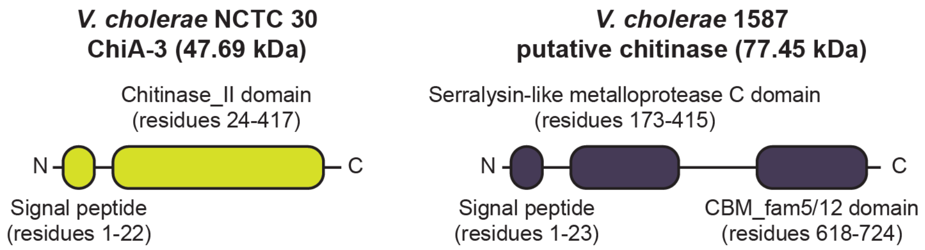


**Supplementary Figure 3. Two putative chitinases are of different sizes and contain different predicted functional domains.** Domain predictions were derived from InterProScan [5] using translated protein sequences obtained from NCTC 30 (gene cluster ‘group_1311’; Figure 2, Supplementary Table 3) and *V. cholerae* 1587 (gene cluster ‘endo_I2’) genome sequences. Images are not to scale.

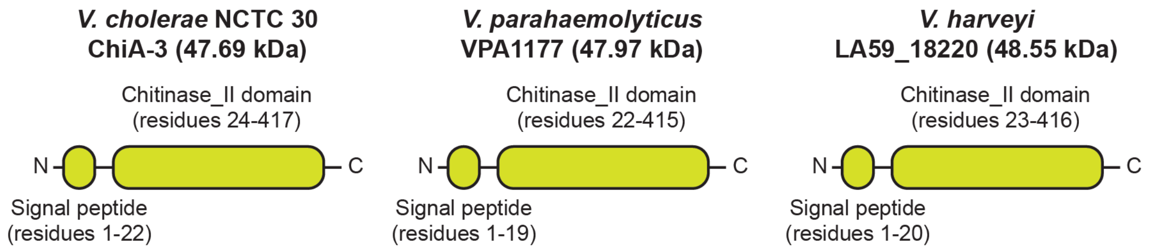


**Supplementary Figure 4. ChiA-3 orthologues from *V. parahaemolyticus* and *V. harveyi* are predicted to have similar domain structures and molecular weights to ChiA-3 from *V. cholerae*.** Domain predictions were derived from InterProScan [5] using protein sequences obtained from the annotated genomes presented in Figure 5b [6, 7]. Images are not to scale.
